# What is the explanation for *Plasmodium vivax* malarial recurrence? Experience of Parasitology Unit of Kinshasa University Hospital of 1982-1983 and 2000-2009

**DOI:** 10.1101/537027

**Authors:** Guyguy Kabundi Tshima

**Author notes:** Correspondence (Guyguy Kabundi Tshima).

## Abstract

**Context and Objectives:** Microscopy is needed for a study involving the surveillance data of a species like *P. vivax*, the most widespread in Asia and almost non-existent species in the Democratic Republic of the Congo (DRC). The use of microscopy and rapid diagnostics tests (RDTs) approaches are recommended for malaria test. Considering the advantages and disadvantages of the two, microscopy is more suitable for effective identification of presence of malaria parasites for the surveillance data of *P. vivax* and other species. Rapid diagnostics tests fit better for *P. falciparum*. This study aimed to revise between the Microscopy and RDTs, which is better for used in city than in rural settings for the surveillance of *Plasmodium vivax* malarial recurrence in malaria-endemic areas and why?

**Methods:** It is a descriptive study of 19,746 laboratory data. The variables wanted were a positive thick drop and a thin smear with the plasmodial species. The analyzes were carried out based on prevalences and the software R was used to generate the figures. The standard threshold of statistical significance was set at 0.05. The ethics committee of the Department of Tropical Medicine approved this study. We were using microscopes as our diagnostic tools for malaria surveillance data in the Parasitology Unit. RDTs are the quickest way to detect and diagnose malaria. It is something that could be easily operated. It can be home-based for everyone depending his understand the principles of how it works. It is also effective and time management. Therefore, this can be used in rural areas because it will be fast to attend to many people. But it has its own limitations because of the differentiation of species. And it not detects *P. vivax*, *P. ovale* and *P. malariae*.

**Results:** From 100% malaria-positive samples, 98.83% were positive for *P. falciparum*, 0.88% were positive for *P. malariae*, 0.063% were positive for *P. ovale*, 0.01% was positive for *P. vivax*. There were co-infections *P. falciparum-P. malariae* representing 0.2%. November 2001 had the high number of positive samples.

**Conclusions:** *P. vivax* at 0.01% highlights that it is an unknown species in the DRC. *P. malariae* at 1% advances our understanding of microscopy utility in the diagnosis of renal failure. *P. falciparum* at 98.83% highlights that it remains the most prevalent species. Efforts for malaria control should be focus on the rain months. Microscopes are effective. Depending on the accurate functionality of the tool and the expertise skill of the technician or scientist. Disadvantages are the facts that it is time consuming. And demands high intellectual understanding of the use of microscopy. Not everyone could operate a microscope. Before you view under the microscope you must prepare the slide and stain to be able to view. All these are long processes. Therefore, microscopy may have lower opportunity to be used in a rural area because of the complexity, the population and time. Microscopy has advantages to be important to use in rural areas because of its accuracy and the ability to detect *Plasmodiums* species than the RDTs.

## Résumé

**Contexte et Objectifs:** La microscopie est nécessaire pour une étude impliquant les données de surveillance d’une espèce comme *P. vivax*, la plus répandue en Asie et une espèce quasi inexistante en République démocratique du Congo (RDC). L’utilisation d’approches de microscopie et de tests de diagnostic rapides est recommandée pour le test du paludisme. Compte tenu de leurs avantages et inconvénients, la microscopie est plus appropriée pour identifier efficacement la présence de parasites du paludisme pour les données de surveillance de *P. vivax* et d’autres espèces. Les tests de diagnostic rapide conviennent mieux à *P. falciparum*. Cette étude visait à réviser entre les tests de microscopie et de diagnostic rapide, ce qui est préférable pour une utilisation en ville qu’en milieu rural pour la surveillance de la récidive du paludisme à *Plasmodium vivax* dans les zones d’endémie palustre et pourquoi?

**Méthodes:** Il s’agit d’une étude descriptive de 19 746 données de laboratoire. Les variables recherchées étaient une goutte épaisse positive et un mince frottis avec l’espèce plasmodiale. Les analyses ont été effectuées sur la base des prévalences et le logiciel R a été utilisé pour générer les chiffres. Le seuil standard de signification statistique a été fixé à 0,05. Le comité d’éthique du Département de médecine tropicale a approuvé cette étude. Nous utilisions des microscopes comme outils de diagnostic des données de surveillance du paludisme dans l’unité de parasitologie. Les tests de diagnostic rapide constituent le moyen le plus rapide de détecter et de diagnostiquer le paludisme. C’est quelque chose qui pourrait être facilement exploité. Cela peut être à la maison pour tout le monde en fonction de sa compréhension des principes de fonctionnement. C’est aussi efficace pour la gestion du temps. Par conséquent, cela peut être utilisé dans les zones rurales car il sera rapide d’assister de nombreuses personnes. Mais les tests de diagnostic rapide ont leurs propres limites en raison de la différenciation des espèces. Et ils ne détectent pas *P. vivax*, *P. ovale* et *P. malariae*.

**Résultats:** Sur 100% échantillons positifs, 98,83% étaient positifs pour *P. falciparum*, 0,88% pour *P. malariae* et 0,063% pour *P. ovale*, 0,01% pour *P. vivax*. Il y avait des co-infections *P. falciparum*-*P. malariae* avec 0,2049%. Novembre 2001 avait le nombre élevé des lames positives.

**Conclusions:** *P. vivax* à 0,01% souligne qu’il s’agit d’une espèce inconnue en RDC. *P. malariae* à 1% fait progresser notre compréhension de l’utilité de la microscopie dans le diagnostic de l’insuffisance rénale. *P. falciparum* à 98,83% souligne qu’il reste l’espèce la plus répandue. Les efforts de lutte contre le paludisme doivent être axés sur les mois de pluie. Les microscopes sont efficaces. En fonction de la fonctionnalité précise de l’outil et de l’expertise du technicien ou du scientifique. Inconvénients-sont les faits que cela prend du temps. Et exige une compréhension intellectuelle élevée de l’utilisation de la microscopie. Tout le monde ne peut pas utiliser un microscope. Avant de regarder sous le microscope, vous devez préparer la lame et la coloration pour pouvoir voir. Tous ces processus sont longs. Par conséquent, la microscopie peut avoir moins de chances d’être utilisée dans une zone rurale en raison de la complexité, de la population et du temps. La microscopie présente des avantages importants en milieu rural en raison de sa précision et de sa capacité à détecter les espèces de Plasmodiums par rapport aux tests de diagnostic rapides.

## Introduction

What is Bio-Distribution (BD)? How and why someone who has any expertise in disease or health can be interested in bio-distribution? Behind this concept, there is the same questioning: what is the pathogenic role of the environment? the distribution of diseases has moved in two directions: medical geography (distribution, prevalence of diseases), and the geography of care with health services (access and use of care) [1].

By BD concept, geographers have found their place next to epidemiologists by delineating populations at risk and giving the environment a high priority [1].

Today, the concept of disease has extended, and it is admitted that the disease is the product of multiple factors and more often the conjunction of genetic and environmental factors [1].

Example 1: African human African trypanosomiasis (HAT), which is a tsetse flies or tsetse fly vector, is found exclusively in southern and western Africa, whereas Chagas disease is found exclusively in South America [1].

Example 2: *Loa loa*, which is exclusively concentrated around the Gulf of Guinea (Central Africa). The displacement of Caucasian or Asian populations towards these zones exposes them to develop these two entities without any geographical constraint [1].

Example 3: *Schistosoma mekongi* and *japonicum schistosomiasis* which are linked to the Asian environment, which are more and more regularly detected in Africa, following the migrations of [1].

Example 4: *Plasmodium vivax*. The populations of central Africa are genetically protected against *Plasmodium vivax* malaria because they are of the Duffy negative blood group (absence of a Duffy receptor favorable to the vivax species: they will however be able to develop this entity Pathological following population migrations or genetic mixing with populations with this gene Duffy Population migrations and some genetic mixing between Asian and African populations will be the basis of the bio-eco-genetic modification certain pathological entities because Asian populations in search of economic investments on the wealth in soil and sub-soil of African almost live in Africa [1].

The focus of this paper was to describe the surveillance of *Plasmodium* species in the Democratic Republic of the Congo (DRC) using Microscopy. How reliable were the data with Microscopy? The paper was focused in the description of *Plasmodium* species in DRC and in the link between months, clinical malaria suspicion and confirmed microscopy because of a need to maintain malaria microscopy expertise diagnosis in DRC with good equipment and personnel financial motivation as rapid diagnosis tests (RDT) is accepted in the hospitals [2]. This study will inspire researchers to favor microscopy over RDT for surveillance of plasmodial species.

### Microscopy

Microscopy is recognized as being the test of reference in the routine diagnosis of malaria, however it is incriminated based on its poor performance in peripheral medical structures reported by some authors, while the performance can be very good in the case where a laboratory has the appropriate equipment and skills which is likely for the Parasitology Unit of the CUK.

Biological confirmation, by microscopy or by rapid diagnostic tests, is essential before the start of treatment, for the following reasons: gradual decrease in the proportion of malaria cases among febrile patients. Ask clinicians to look for and treat the true cause of fever for patients who test negative for malaria. The second reason is to avoid wasting antimalarials especially artemisinin derivatives and thus reduce the selective pressure to delay the spread of resistance [3].

The diagnosis of malaria is a relevant and dynamic topic, with many publications [3].

The Global Technical Strategy for Malaria 2016-2030, endorsed by the World Health Assembly in May 2015, sets ambitious and achievable targets for 2030, including reducing the incidence of malaria by at least 90% and associated mortality [3], which requires good diagnostic tools with competent staff.

Unfortunately, 86% of all malaria cases are recruited in sub-Saharan Africa and *A. gambiae* seems to be the most implicated vector in which *P. falciparum* is present in more than 90% of cases. It is also responsible for severe forms of the disease [3].

The DRC is still a major endemic country for malaria just after Nigeria. In the world and despite the progress made, 40% of malaria deaths are still in the two countries mentioned above [2].

### Specific objectives

- Determine the number of samples taken in the study period 2000-2009
- Identify the proportion of positive results per year
- Estimate the parasitic density per year
- Evaluate the distribution of *Plasmodium* species
- Determine the proportion of positive cases per month with high confirmed samples

## Material and methods

### Study zone

This study was conducted at the Unit of Parasitology of the CUK. This service has a parasitology reference laboratory. The expertise and the equipment of this laboratory make possible to measure the malaria parasite density but also to identify the species of *Plasmodium* in the blood.

### Design of the study

The data were collected descriptively from all patients who visited the CUK and who were referred to the Parasitology Unit for Malaria Screening from 2000 to 2009. Microscopy was used to identify the different plasmodial species in blood samples obtained from 19,746 patients for suspected clinical malaria.

### Methods

The analysis of the data obtained from the laboratory data of the Parasitology Unit served as a data source. In this work, I describe the results of microscopy for the diagnosis of malaria for 10 years. The classic threshold of statistical significance is 0.05 in our calculations.

### Study procedure

The participants were selected basis on the clinical diagnosis of malaria which was done according to physical signs: This included clinical history, vital signs, body weight, signs of dehydration, and neurological and physical examinations (skin, abdominal, ear, mouth, throat). Clinical examination, socio-demographic information and Medical history were performed on each patient by the CUK clinicians.

### Statistical analysis

Extracted data were managed in a Microsoft Excel database and analyzed descriptively on four tables. All statistical computations were performed in R for drawing figures. The relationship between microscopy and seasons characteristics and the presence of malaria parasites was assessed via R. The total counts of Microscopy tests were used to calculate the prevalence. Only the baseline data collected from 2000 to 2009 is presented.

### Codification used for graphs

#Figure 1

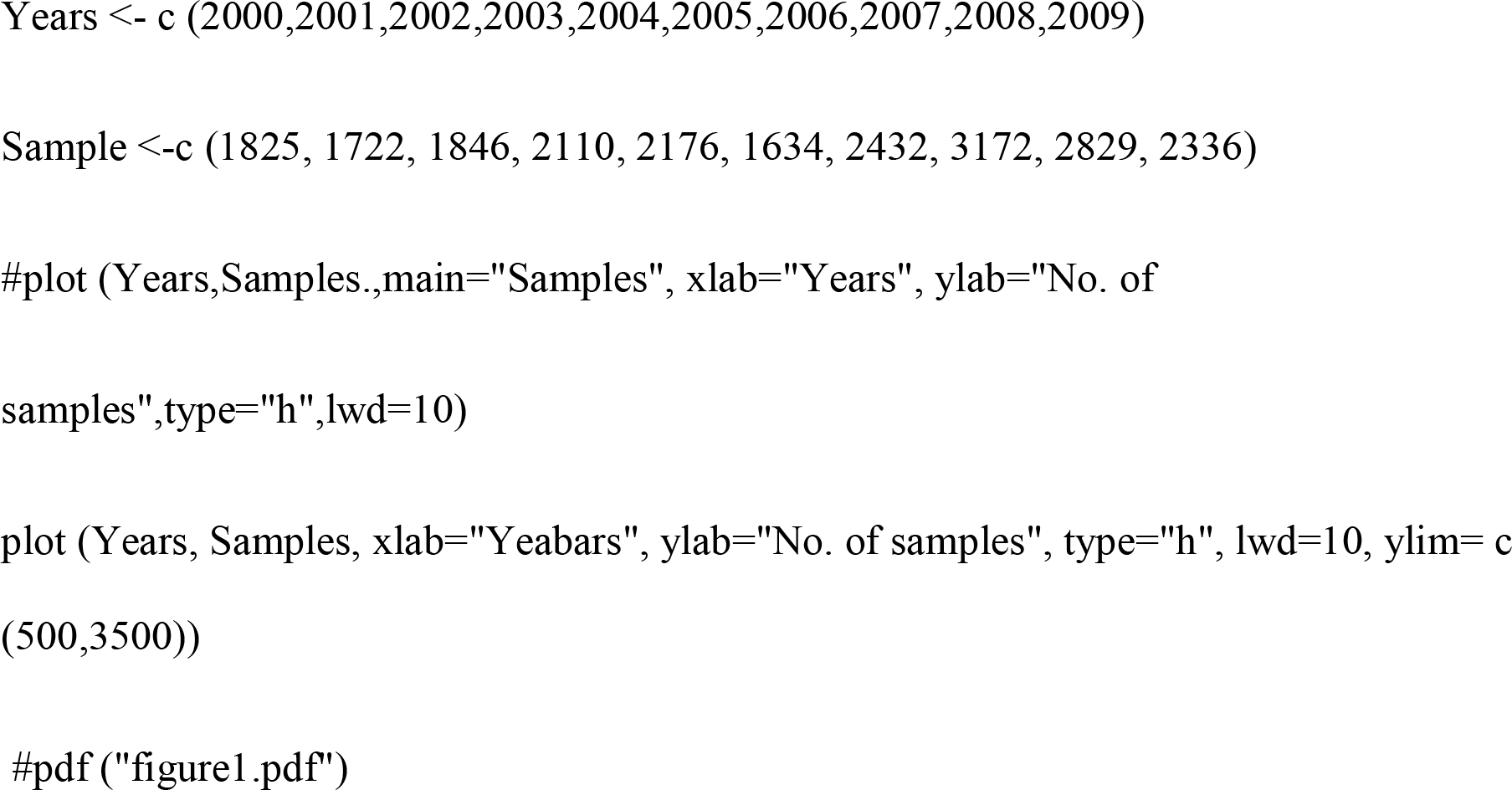

#Figure 2

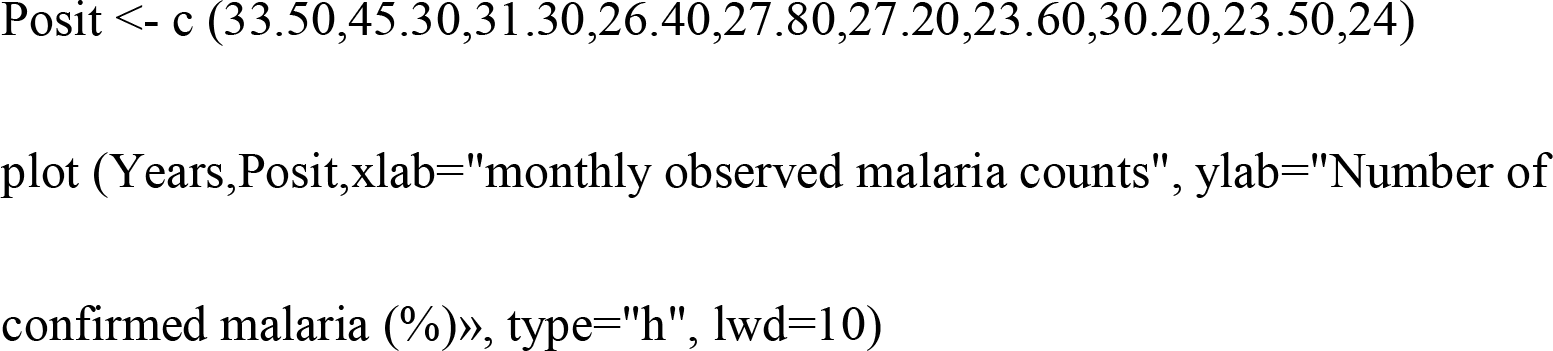

#Figure 3

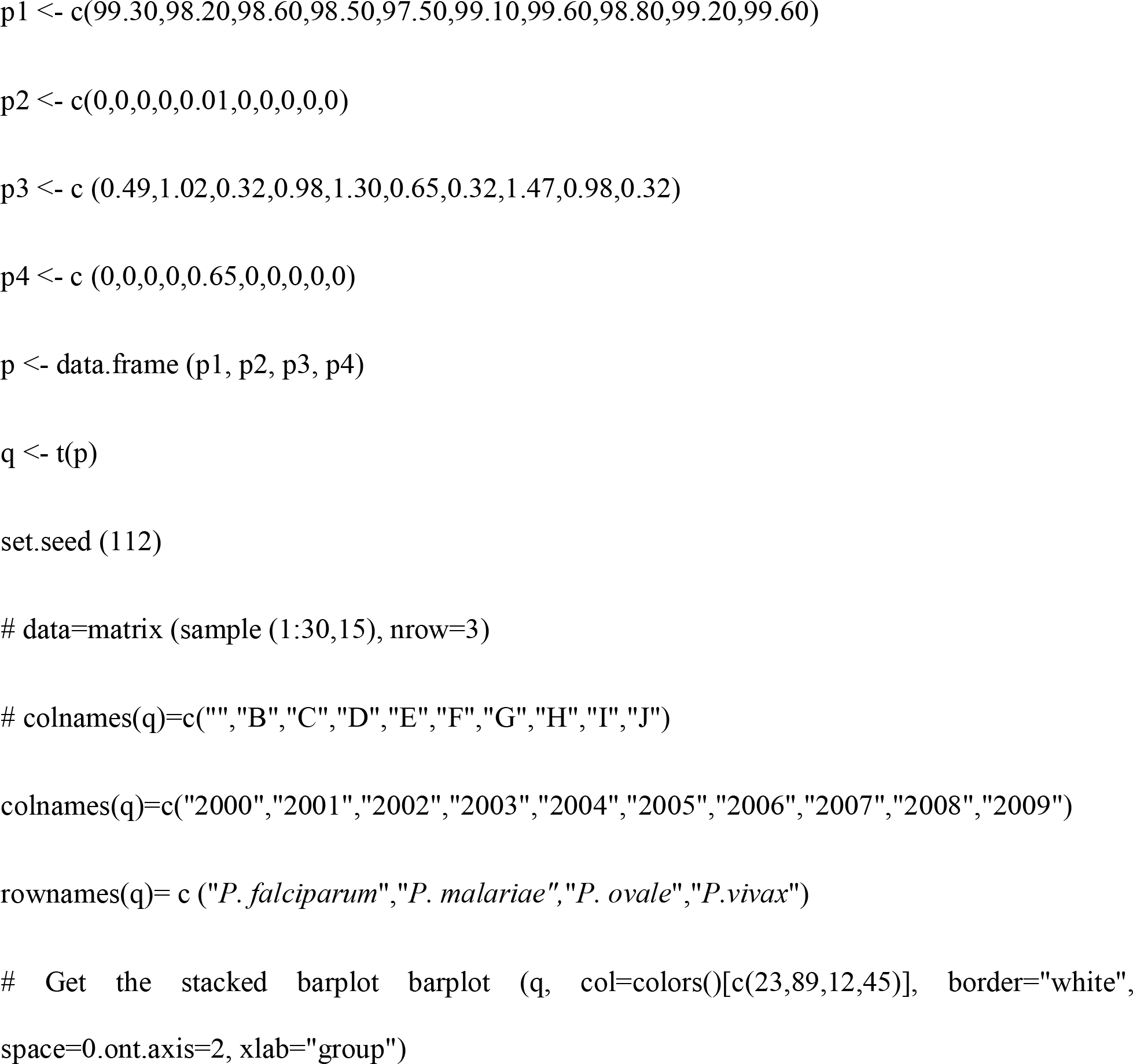

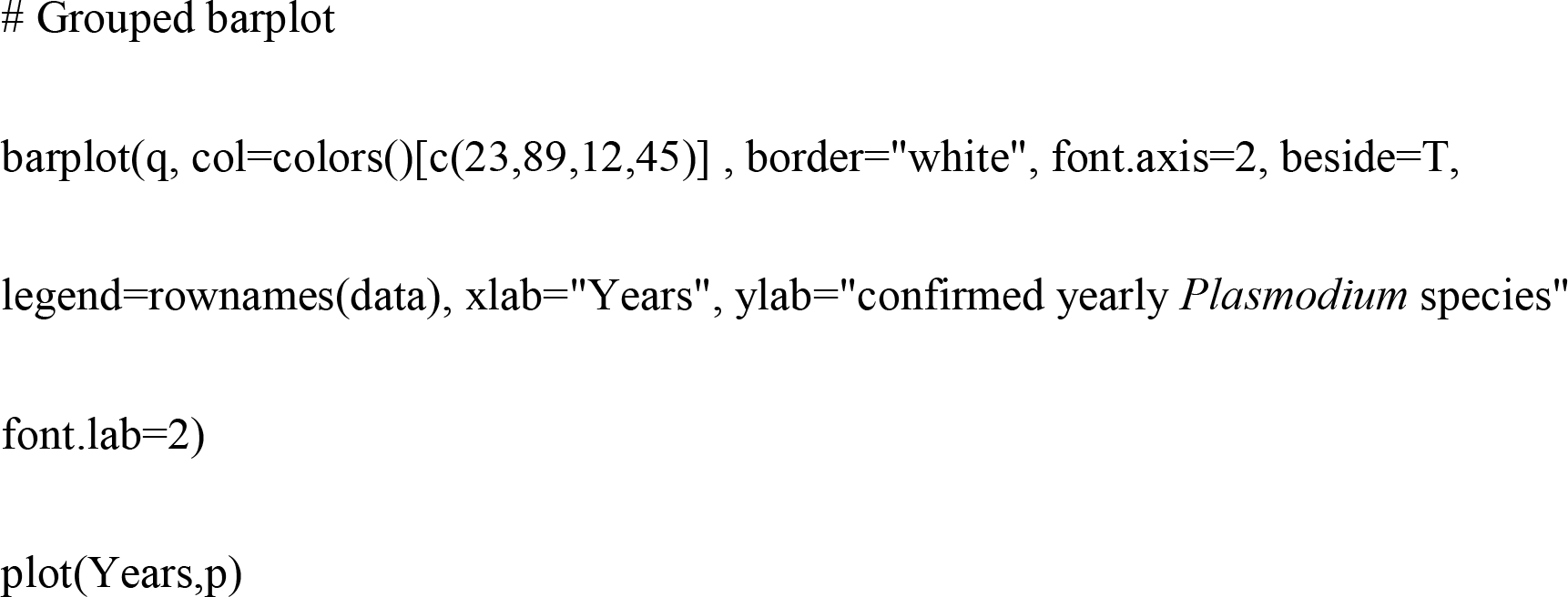

#Figure 4

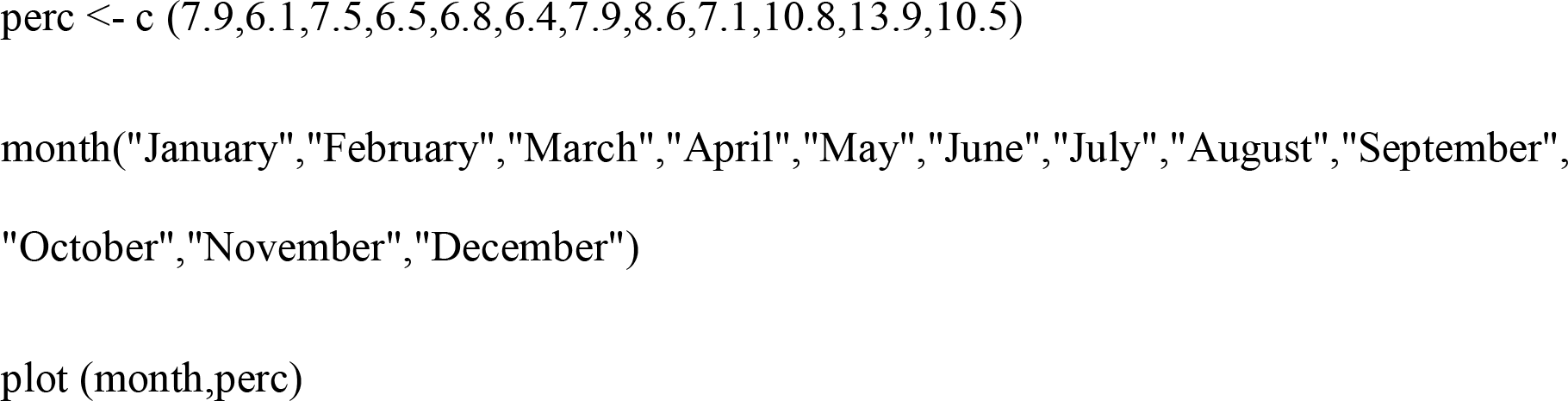

#Figure 5

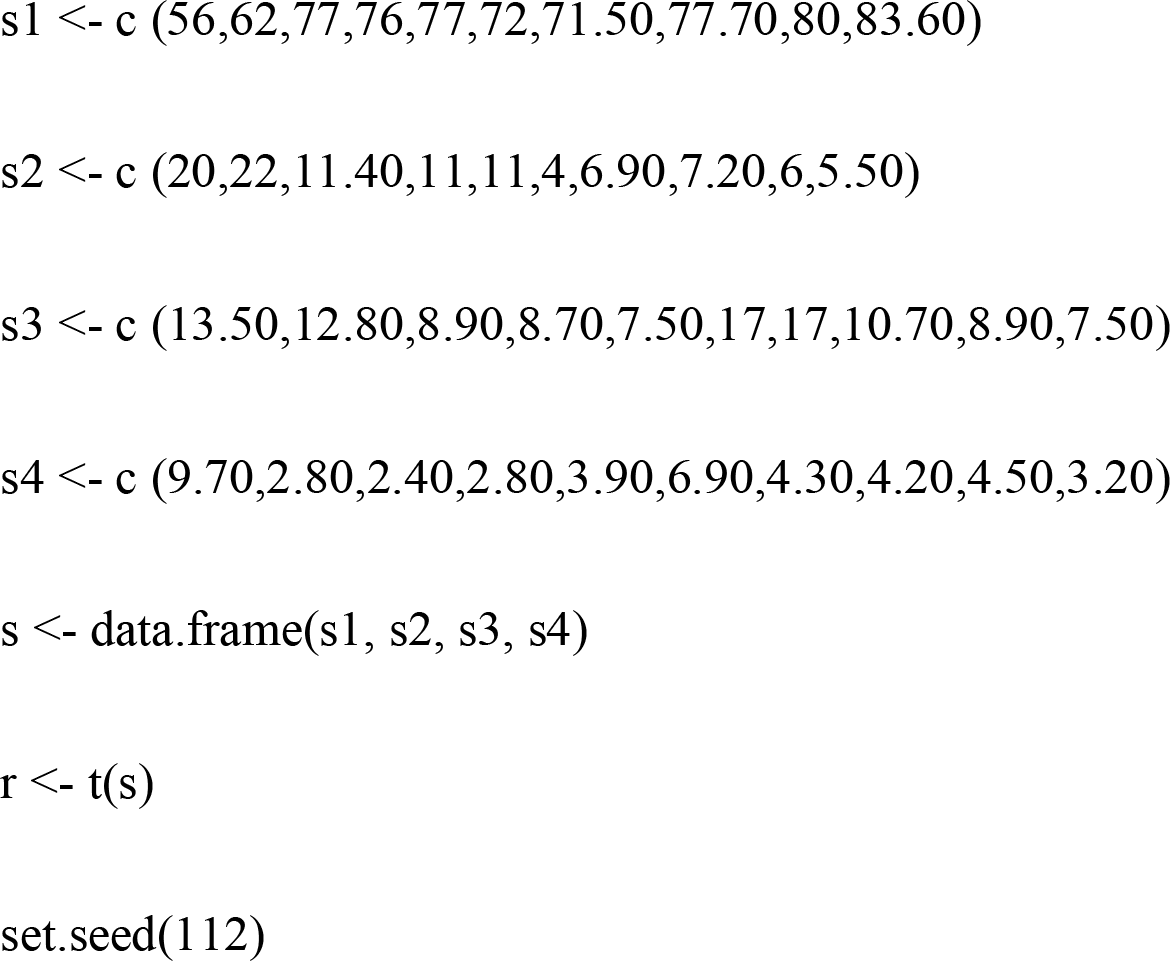

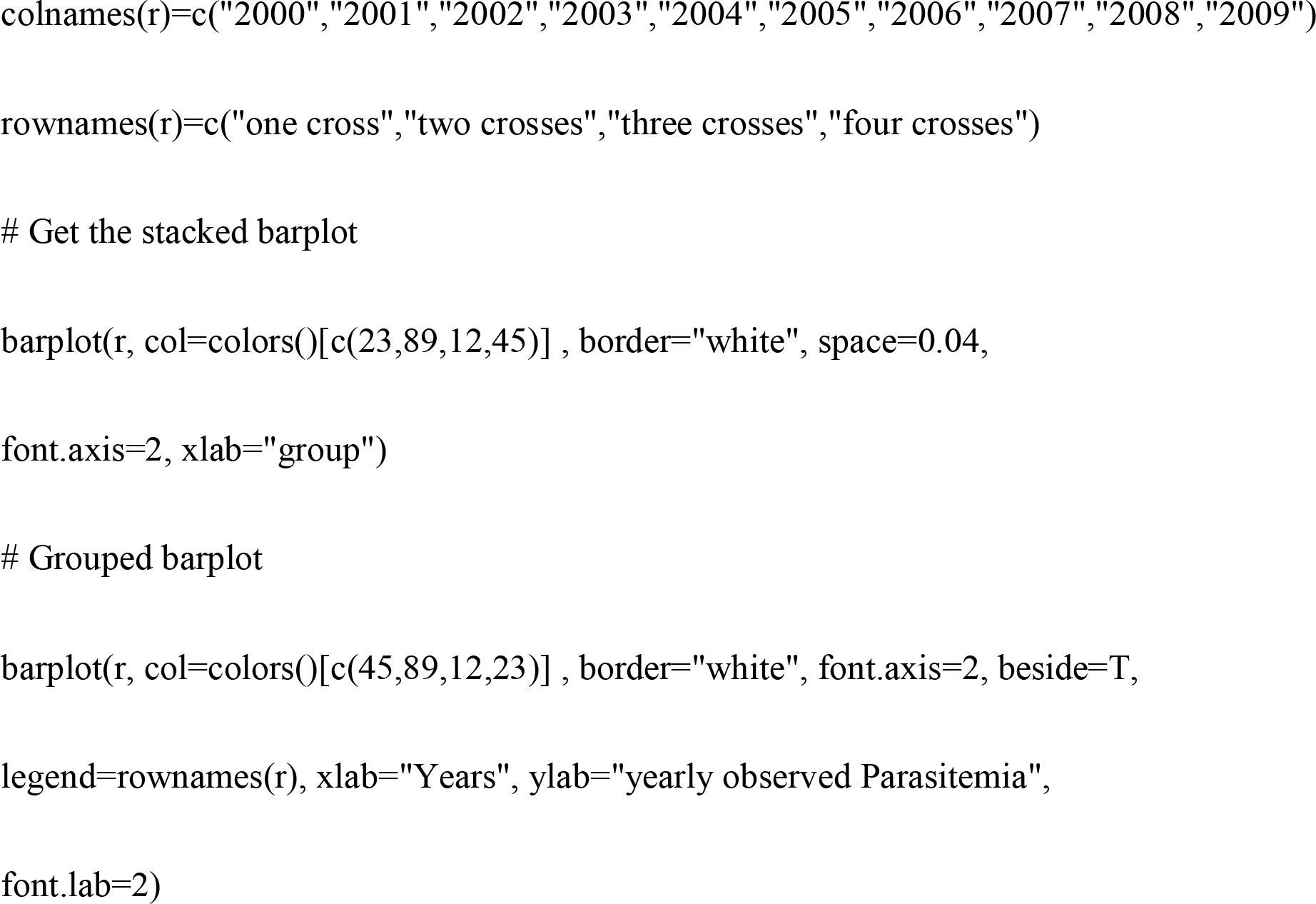

## Results

Number of samples taken in the study period of 10 years:

The present work examined the results of the parasitological analyzes of malaria carried out at the Parasitology Unit of the Kinshasa University Hospital in a period of 10 years. I collected a total of 19,746 samples. And 6,344 samples were positive, or 32% (Table 1).

**Table 1.** The total yearly observed malaria burden from 2000-2009 in Kinshasa

| Year | No. of samples examined | Confirmed malaria | Percentage |
| --- | --- | --- | --- |
| 2000 | 1825 | 612 | 33.50% |
| 2001 | 1722 | 781 | 45.30% |
| 2002 | 1846 | 578 | 31.30% |
| 2003 | 2110 | 558 | 26.40% |
| 2004 | 2176 | 609 | 27.80% |
| 2005 | 1634 | 446 | 27.20% |
| 2006 | 2432 | 574 | 23.60% |
| 2007 | 3172 | 959 | 30.20% |
| 2008 | 2829 | 666 | 23.50% |
| 2009 | 2336 | 561 | 24% |
| Total | 19746 | 6344 | 32% |

### Proportion of high demands per year

The year 2007 was marked by the highest number of requests for laboratory tests for the diagnosis of malaria in Parasitology Unit (Table 1). Curiously, the frequency of positive cases was lower than in 2001 (30.2% vs. 45.3%) (Table 1).

### Proportion of positive results per year

The data show that the highest number of positive cases was recorded in 2001: 45.03% (Table 1)

### Proportion of positive results per month in 2001

Table 2 showed that malaria is a seasonal disease depending on rain months divided in two periods: heavy rain period from October to December and low rain period from January to March.

**Table 2.** The monthly confirmed malaria counts observed in 2001

| Study period | Number of samples | Percentage |
| --- | --- | --- |
| January | 136 | 7.9 |
| February | 105 | 6.1 |
| March | 129 | 7.5 |
| April | 113 | 6.5 |
| May | 117 | 6.8 |
| June | 110 | 6.4 |
| July | 136 | 7.9 |
| August | 148 | 8.6 |
| September | 122 | 7.1 |
| October | 186 | 10.8 |
| November | 239 | 13.9 |
| December | 181 | 10.5 |
| Total | 1722 | 100% |

### Parasitemia

The data in table 3 indicate that most patients had parasitaemia at a cross or low parasitaemia.

**Table 3.** Observed Yearly parasitaemia

| Year | One cross | Two crosses | Three crosses | Four crosses |
| --- | --- | --- | --- | --- |
| 2000 | 342/612 | 122/612 | 82/612 | 59/612 |
| 2001 | 484/781 | 172/781 | 100/781 | 22/781 |
| 2002 | 445/578 | 66/578 | 51/578 | 14/578 |
| 2003 | 424/558 | 61/558 | 49/558 | 16/558 |
| 2004 | 469/609 | 67/609 | 46/609 | 12/609 |
| 2005 | 321/446 | 18/446 | 76/446 | 31/446 |
| 2006 | 410/574 | 40/574 | 98/574 | 25/574 |
| 2007 | 745/959 | 69/959 | 103/959 | 40/959 |
| 2008 | 533/666 | 40/666 | 59/666 | 30/666 |
| 2009 | 469/561 | 31/561 | 42/561 | 18/561 |
| Total | 4642/6344 | 686/6344 | 706/6344 | 267/6344 |

### Distribution of Plasmodium species

From 100% malaria-positive samples, 98.83% were positive for *P. falciparum*, 0.88% were positive for *P. malariae*, 0.063% were positive for *P. ovale*, 0.01% was positive for *P. vivax*. There were co-infections *P. falciparum- P. malariae* representing 0.2% (Table 4). The presence of *P. vivax*, but with only 1 case seems interesting (Table 4).

**Table 4.** Annual distribution of *Plasmodium* species from 2000 to 2009 in Kinshasa

| Year | Samples | Confirmed | *P.falciparum* | *P.malariae* | *P.ovale* | *P.vivax* | Co-infections *P.falciparum*/*P.malariae* |
| --- | --- | --- | --- | --- | --- | --- | --- |
| 2000 | 1825 | 612 | 608 | 3 | 0 | 0 | 1 |
| 2001 | 1722 | 781 | 767 | 8 | 0 | 0 | 0 |
| 2002 | 1846 | 578 | 570 | 2 | 0 | 0 | 6 |
| 2003 | 2110 | 558 | 550 | 6 | 0 | 0 | 2 |
| 2004 | 2176 | 609 | 594 | 8 | 4 | 1 | 2 |
| 2005 | 1634 | 446 | 442 | 4 | 0 | 0 | 0 |
| 2006 | 2432 | 574 | 572 | 2 | 0 | 0 | 0 |
| 2007 | 3172 | 959 | 948 | 9 | 0 | 0 | 2 |
| 2008 | 2829 | 666 | 660 | 6 | 0 | 0 | 0 |
| 2009 | 2336 | 561 | 559 | 2 | 0 | 0 | 0 |
| Total | 19746 | 6344 | 6270 | 50 | 4 | 1 | 13 |

The rest of the results are shown in Figures 1 to 5: The data in Figure 1 indicate that the highest demand for parasitological examinations occurred in 2007. The data in Figure 2 show that the highest number of positive cases was recorded in 2001.The data in Figure 3 show that the month of November was marked by the largest number of positive cases. The data in Figure 4 reveal that most patients had parasitaemia on a cross. The data in Figure 5 indicate that *P. falciparum* was the most isolated plasmodial species.

**Figure 1.**
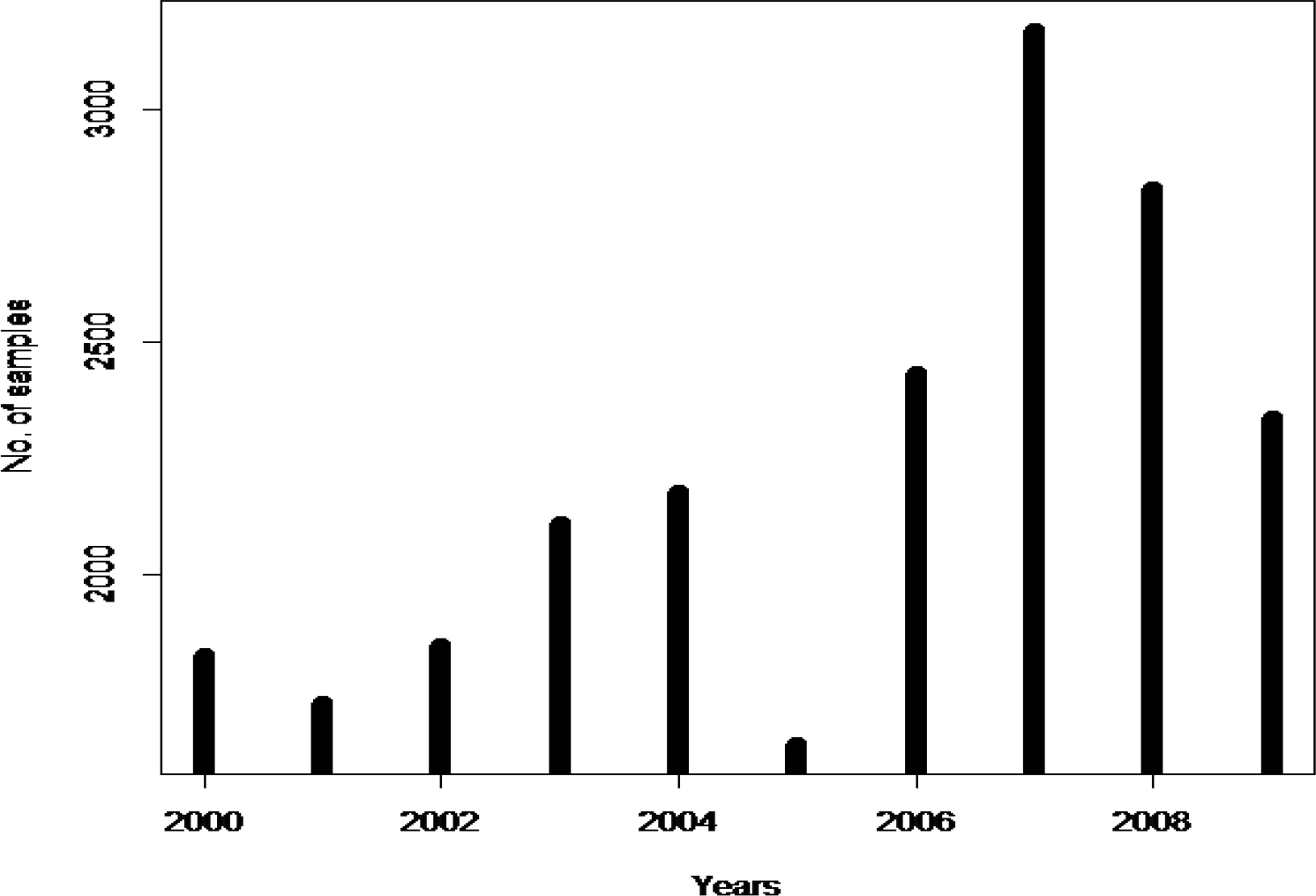
Histogram of high demands per year. The above data indicate that the highest demand for parasitological examinations occurred in 2007.

**Figure 2.**
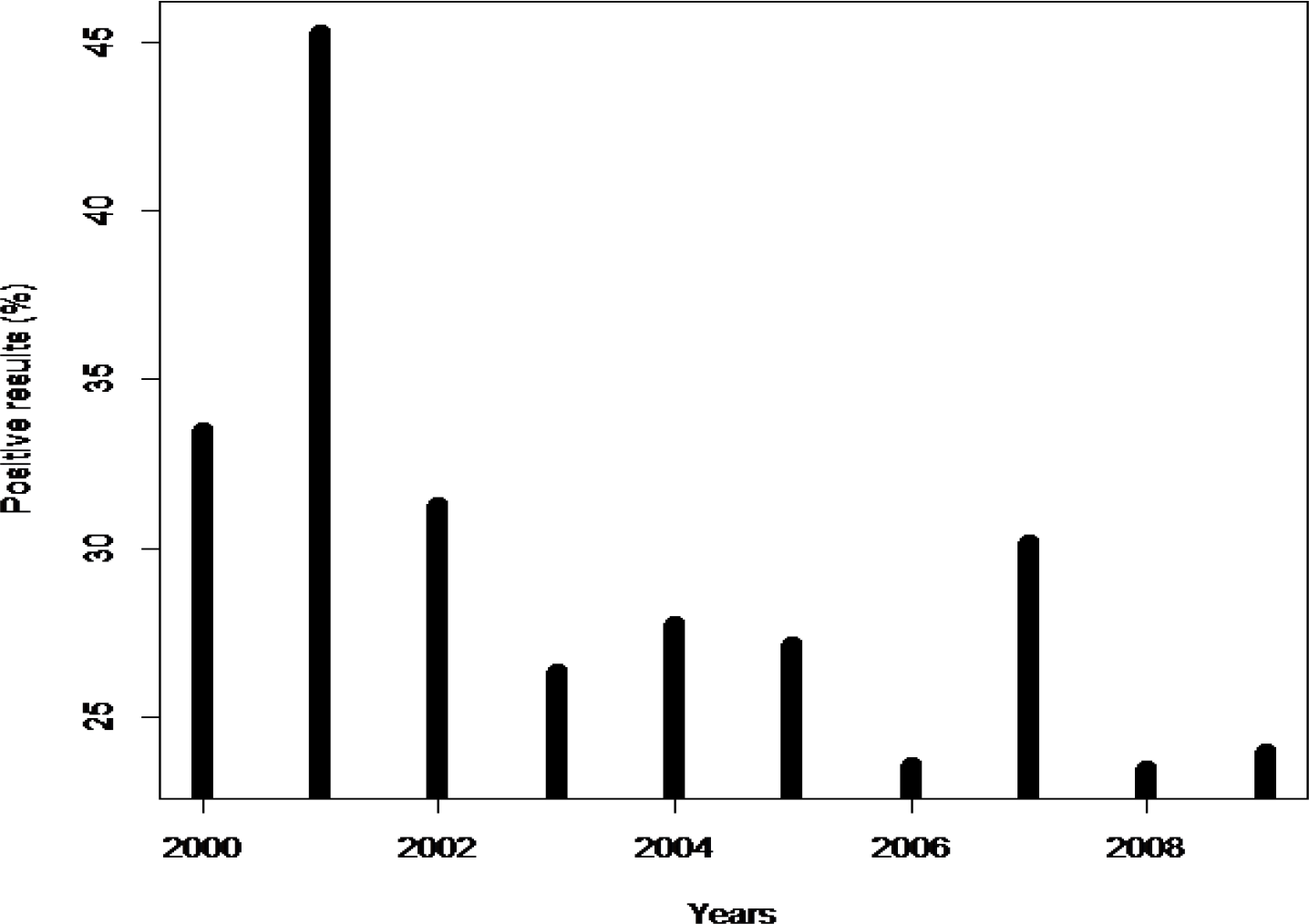
Histogram of high confirmed malaria counts per year. The data show that the highest number of positive cases was recorded in 2001, 45.03% (p <0.05).

**Figure 3.**
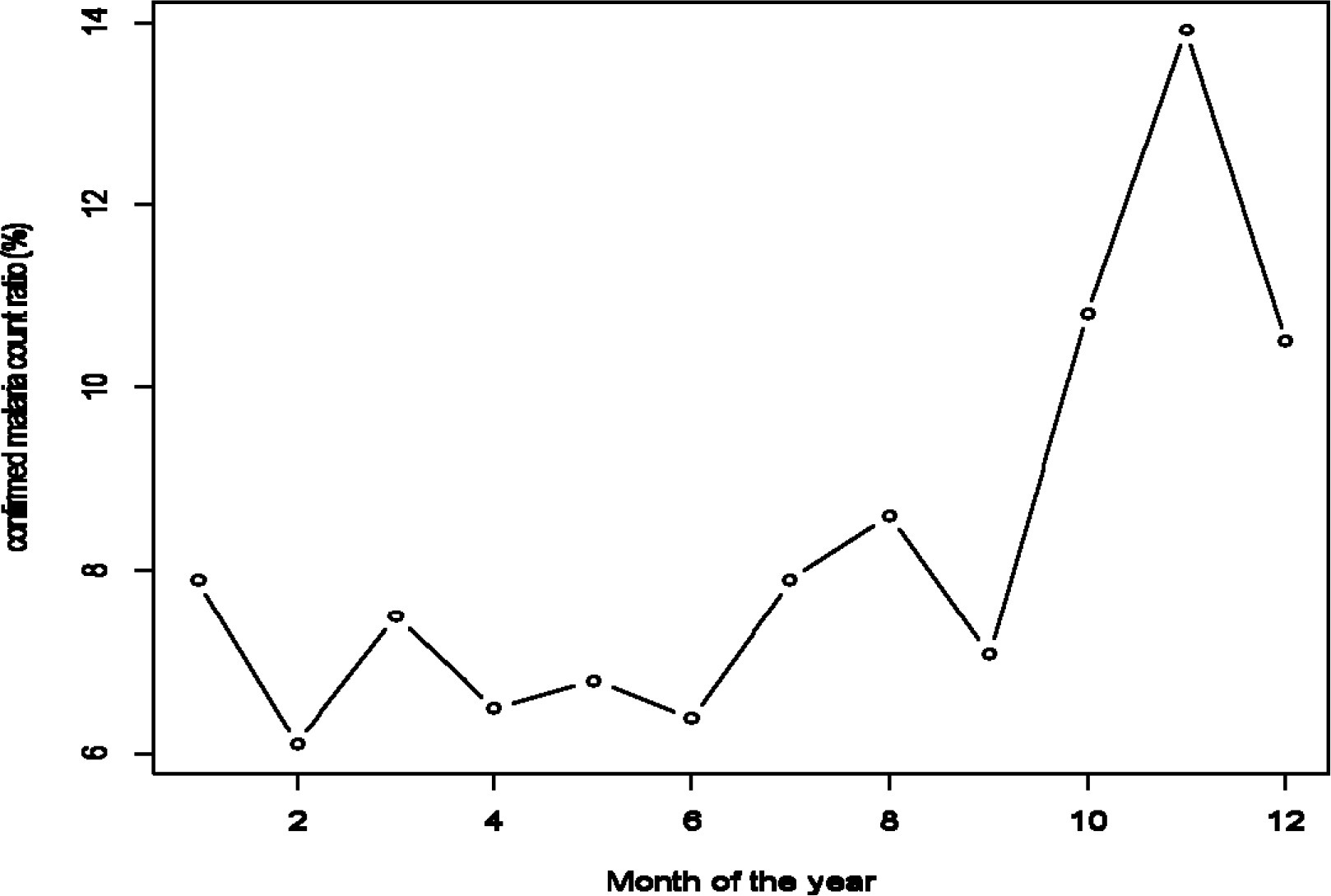
Plot chart of confirmed malaria counts per month in 2001. The month of November was marked by the largest number of positive cases.

**Figure 4.**
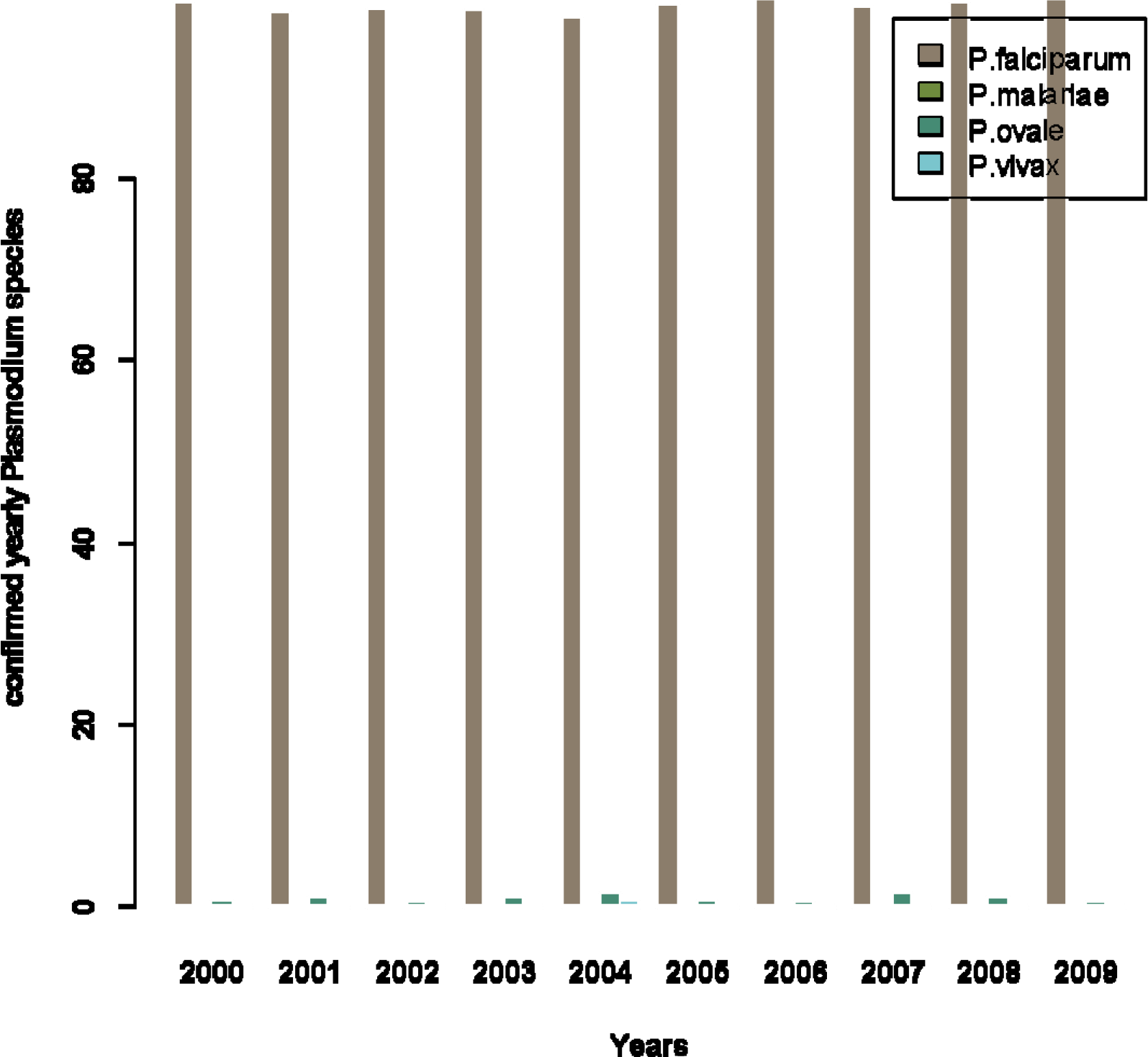
The total of confirmed yearly *Plasmodium species*. The data indicate that *P. falciparum* was the most isolated plasmodial species.

**Figure 5.**
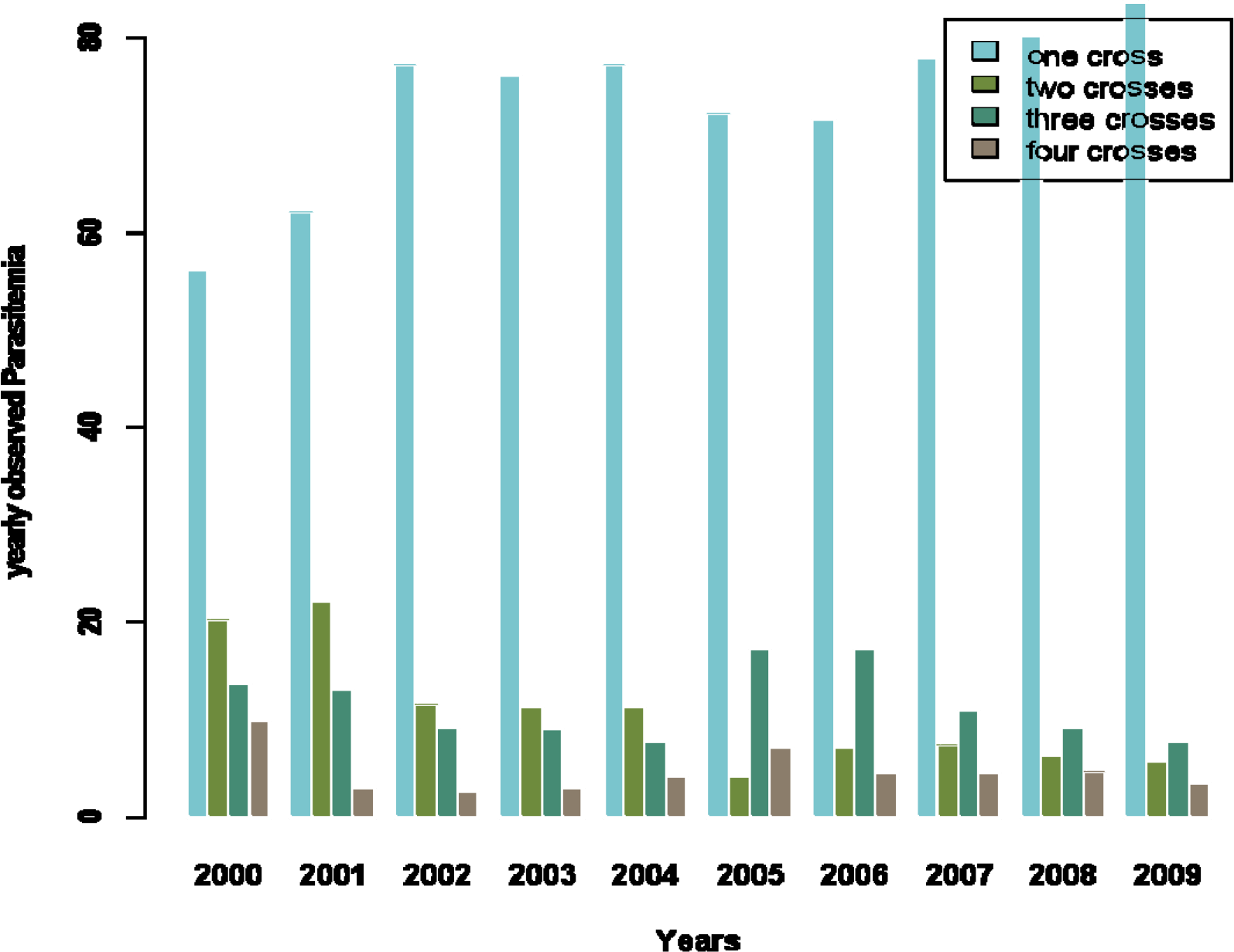
Plot of the total observed yearly *Parasitemia*. Parasitaemia (Parasitic density per year): These data reveal that most patients had parasitaemia on a cross.

## Discussion

The present study focused on the determination of the biodistribution of *Plasmodium* species in Kinshasa, with the reference laboratory of the Unit of Parasitology of the Department of Tropical Medicine, Infectious and Parasitic Diseases of the Kinshasa University Hospital as source of information. The data collected covers the period from 2000 to 2009, a period of 10 years.

### Number of samples taken in the study period of 10 years

Of 19,746 microscopy samples tested over 10 years; 6,344 were found positive, namely 32%. These data confirm those of the literature [4, 5]: Ngimbi et al., in their Kinshasa study on malaria parasitaemia in 1982, reported a frequency of 33.8% [4]. In 1984, they reported a frequency of 33.2% [5]. Both studies corroborate our results.

### Proportion of high demands per year

The number of positive cases was recorded in 2001. 2001 was marked by the beginning of the resistance on antimalarials drugs involving a change towards the artemisinin derivatives, but it was in 2005 that the national malaria control programme PNLP introduced the combination of artesunate-amodiaquine to treat cases of uncomplicated malaria or simple malaria forms, also Artemether-Lumefantrine and Dihydroartemisinin-piperaquine for complicate malaria forms. The old combination was sulfadoxine and pyrimethamine for uncomplicate malaria and quinine for complicate malaria forms [2].

### Proportion of high confirmed malaria counts per year

The results of this study show that 32% (6,344 / 19,746) samples were positive. We recommend the use of biological diagnosis before treating patients. In general, this work highlights the reflection on the concerns of developing effective tools for the early diagnosis of infectious agents, particularly in generally poor endemic areas, such as Kinshasa, capital of the DRC. Automated diagnosis uses the detection of plasmodial pigment with PCR is more helpful because of its high sensitivity and specificity. Without powerful tools at low cost, the therapeutic test is the last alternative in the search for solutions to better treat patients and obviously, it does not avoid unnecessary treatments and damage the Congolese family income already poorer. Our reminder is consistent with the logic of associating the laboratory with the clinic. Several new methods of diagnosing malaria have recently been developed, but all are based on clinical suspicion and, therefore, an explicit clinical demand is required. Without clinician demand for regular malaria screening, the malaria diagnosis may be missed, and it is desirable to proceed to clinical automated screening with some methods that lend themselves to automation (e.g. PCR) with a new generation of tools. Comprehensive blood count analyzers, widely used in clinical laboratories, and which have the potential to detect hemozoin in white blood cells and possibly in erythrocytes.

### Proportion of confirmed malaria counts per month in 2001

The highest parasitaemia was observed in 2001; It was also observed in the last quarter of the year with a pic or the highest number of confirmed samples at the month of November (Figure 5).

In Kinshasa, the last three months of the year is the period of heavy rain with temperatures between 30 ° C and 38 ° C [6]. These conditions are favorable to the proliferation of *Anopheles* that would promote the transmission of malaria during this time of the year without the use of mosquito measures.

Our results reinforce the study conducted by Mulumba et al., they clearly contrasted the epidemiological aspects of the rainy periods (March-May and October-December) with dry periods (January-February and June-September) [6].

Mulumba showed a high prevalence of malaria during the rainy season compared to the dry season [6]. Thus, the change of seasons (long rainy season to short dry season) is a risk factor that must be considered in improving malaria control strategies [6].

### Parasitaemia

The data in Figure 2 indicate that most patients had parasitaemia at a cross or low parasitaemia. Preimmunization reduces the multiplication of *Plasmodium* in the body of the host. This would explain the low parasitaemia. It’s an advantage of pre-immunization that protect subjects to suffer of severe and complicated forms of malaria without immunodeficiency. With immunodeficiency, malaria is serious.

Particular attention should be paid to migrants non-immune in the era of globalization. Four categories of migrants at risk of malaria have just been described in the DRC: Subjects coming from a malaria free area; Subjects who has left the DRC for more than 6 months and who returns; Subjects from an unstable transmission malaria area (mountain facies) to a stable transmission zone (tropical / equatorial facies); Subjects expatriate in precarious situations because they have to live far from home [2]. The migrants fall into the category of YOPI group: Young, Old, Pregnants, Immigrants and Immunodeficients. The YOPI group is vulnerable to severe forms of malaria. Thus, a preventive treatment is reserved for this category: For Young and Old, whose stay does not exceed 6 months, Atovaquone-Proguanil (Malarone®) may be prescribed [2].

For Pregnants women: Intermittent Preventive Therapy is Sulfadoxine-Pyrimethamine [2];

### Distribution of Plasmodium species

For a long time, it has been recognized that *P. vivax* has not been present in countries where the population does not have Duffy antigen. That is the situation in the DRC. Its isolation in the blood of a Congolese would be the result of mixed marriages with other nationalities who have Duffy antigen. In addition, it should be noted that some provinces of the DRC share common borders with countries where this species of *Plasmodium* occurs. This is especially true in the Province Orientale.

However, I worried about the frequency of *P. malariae* even with low percentage because of its involvement especially in the development of nephrotic syndrome with renal failure.

About literature data on *Plasmodiums* species, Ngimbi et al. in their study in Kinshasa in 1982, reported the following frequencies: 97.9% for *P. falciparum*, *P. malariae* 0.9%, 0.2% for *P. vivax* and 0.6% for *P. ovale* [4]. *P. malariae*, *P. vivax* and *P. ovale* showed a lower frequency in the present work.

This difference is because Ngimbi et al. had worked in the community (Livulu, Kisenso, Ngaba, Barumbu, Kimbanseke, Mount Ngafula) and they had received patients with the first symptoms of the disease in the health centers [4], while my data came from the laboratory of a tertiary level hospital where patients come for last-resort consultation after visiting polyclinics and other health centers without success.

*Plasmodium falciparum* was the most found plasmodial species in this work. However, the persistence of *P. falciparum* represents a real threat. In fact, it is responsible for severe and complicated malaria access. It is involved in most of the problems of antimalarial drug resistance. *P.falciparum* at 98.83% highlights that it remains the most prevalent to inform control and elimination strategies as it highlighted by these authors [7].

## Conclusion

*P. vivax* at 0.01% highlights that it is an unknown species in the DRC. Microscopy combined with automated systems will be an answer to the thorny issue of diagnosis of *Plasmodium* species in the laboratory.

Based on the data in this study, we recommend that the Parasitology Unit establish a formal collaboration with the PNLP in order to update *Plasmodium* map and adapt control strategies. Closer collaboration with other reference laboratories in the country in multi-center studies and inter-laboratory quality control is warmly welcome.

## Acknowledgments

None

## References

1. Wumba R. Géographie de la santé à la croisée de la géographie des maladies et de la géographie de soins. Ann. Afr. Med., 2018: 2

2. Situakibanza NTH. Quoi de neuf dans la lutte contre le paludisme en République Démocratique du Congo depuis 2014? Ann. Afr. Med., 2016: 9

3. Rapport mondial du paludisme (OMS 2015).

4. Ngimbi NP, Beckers A, Wery M. Aperçu de la situation épidémiologique du paludisme à Kinshasa (République du Zaïre) en 1980. Ann Soc belge Med Trop 1982; 62 : 121–37.

5. Ngimbi NP, Wéry M, Henry MC, Mulumba MP. Réponse in vivo à la chloroquine au cours du traitement du paludisme à Plasmodium falciparum en région suburbaine de Kinshasa, Zaïre. Ann Soc belge Med Trop 1985; 65 (Suppl 2) : 123–35.

6. Mulumba MP, Muyembe TJJ. Diagnostic de l’infection palustre à Kinshasa: contrôle de qualité inter et intralaboratoire. Congo Médical 1999; 2 :759–69.

7. Lindsey Wu, Lotus L. van den Hoogen, Hannah Slater, Patrick G. T. Walker, Azra C. Ghani, Chris J. Drakeley & Lucy C. Okell. Comparison of diagnostics for the detection of asymptomatic Plasmodium falciparum infections to inform control and elimination strategies. Nature 2015; 528, S86–S93, doi: 10.1038/nature16039.

